# Bayesian phylogenetic analysis for diploid and allotetraploid species networks

**DOI:** 10.1101/129361

**Authors:** Graham Jones

## Abstract

Allopolyploid species are formed by genome doubling after hybridization between otherwise intersterile parental species. Allopolyploidy is a common speciation mechanism in land plants. Here we describe and evaluate a Bayesian approach to the phylogenetic analysis of species relationships when both ordinary speciation and allopolyploidy are present. The approach takes incomplete lineage sorting into account using the multi-species coalescent model, and extends this to deal with the extra complications due to allopolyploidy. The number of hybridizations is not assumed, which means that the number of parameters varies and a reversible-jump MCMC algorithm is needed to sample from the posterior. The main restriction is that only diploids and allotetraploids are considered. The model is implemented in the BEAST framework and is an extension of Jones et al. (2013). Simulations show that the topology of the network can be reliably inferred along with estimates of other parameters.

## 1 Introduction

The aim of this paper is to describe and evaluate a Bayesian approach to the phylogenetic analysis of species relationships when both ordinary speciation and allopolyploidy are present. It builds on the work in Jones et al. (2013) which was restricted to a single hybridization. The main restriction here is that only diploid and allotetraploid species are considered. Thus we assume that the species being analyzed have undergone at most one round of allopolyploidization since the root of the species network. We also assume that within the allotetraploids, there is no recombination between sequences from different parental species. This means that all the sequences can be seen as the result of the evolution of diploid genomes, but after hybridization, the topology, node times, and population sizes are shared.

The evolutionary history can be represented as a network or as a multi-labeled tree (MUL-tree), which is a binary tree in which more than one tip may be labeled by the same species. Both these representations omit information about extinct species; they are reconstructions from extant taxa. An example is shown in Figure 1. This shows a scenario which results in two extant diploid species a and b, and three extant allotetraploid species x, y, and z. Reading these diagrams from left to right, there are three ordinary speciations resulting in four diploid species before the first hybridization. Then two of these hybridize to form allotetraploid species x which continues to present. This is followed by two more ordinary speciations of diploid species, and a second hybridization to produce an allotetraploid species, which then undergoes ordinary speciation to produce y and z. The diploid species which contribute to the hybridizations become extinct before present time, leaving a and b. Note that it is straightforward to convert the network representation into a unique multi-labeled tree representation. There are algorithms for the reverse operation (Huber and Moulton, 2006) but the network obtained is not in general unique.

**Figure 1:**
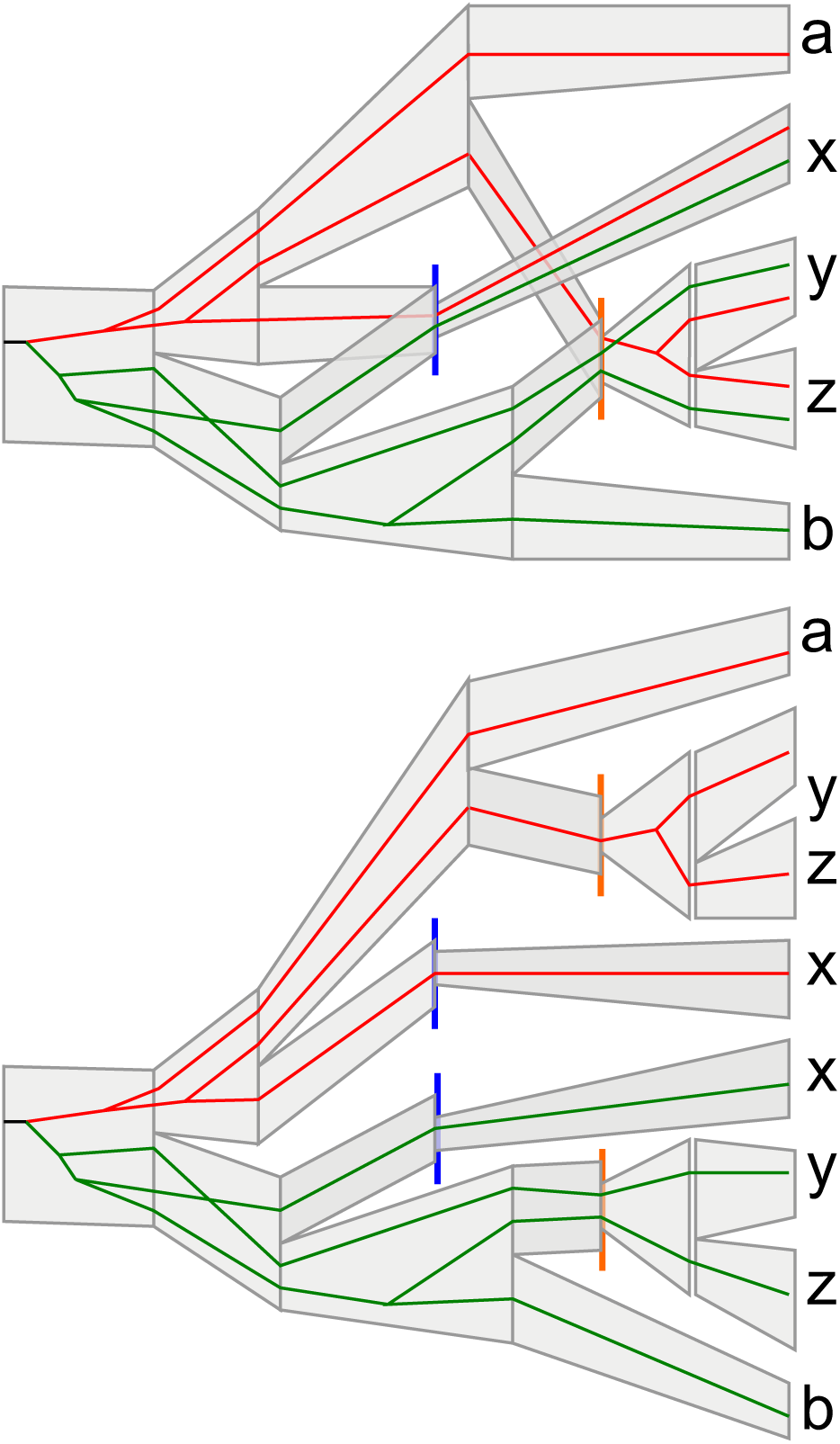
Top: a species network for two diploid species a and b, and three allotetraploid species x, y, and z. The widths of the gray ‘tubes’ indicate population sizes. The network contains a gene tree, with red and green branches indicating different parental species. The blue line indicates one hybridization, the orange lines another. Bottom: the same scenario represented as a multi-labeled tree.

There are several problems to deal with in the phylogenetic analysis. In common with the inference of species trees in which there is only ordinary speciation, both mutational variance (due to the stochasticity of mutations) and coalescent variance (due to the stochasticity of mating) are present. Coalescent variance can produce incomplete lineage sorting. Thus we simultaneously estimate the gene trees and the species network into which they fit. We use a generalization of the multispecies coalescent model to deal with this. Secondly, when the DNA from allotetraploid organisms is sequenced, it is not possible a priori to assign sequences to their parental diploid species. Thus there is an ambiguity in the labeling of the sequences which is not normally present. These two issues were dealt with in Jones et al. (2013), but there only a single hybridization was assumed. Here we also infer the number of hybridizations. This means that inference must explore a space of species networks in which the number of parameters (node times and population size parameters) varies. We use a reversible-jump MCMC process to explore this space.

The method is implemented in BEAST 1.8 and is called AlloppNET. A manual and R scripts are available from the author’s web site (

~~~
http://indriid.com/workingnotes2013.html
~~~

) to help generate the BEAST XML file. We evaluate the method on simulated data only in this paper. Rothfels et al. (2017) have used AlloppNET to estimate a network for 9 diploid species and 19 tetraploid species from the fern family *Cystopteridaceae*.

## 2 Model and priors

There is a large amount of notation which we collect here. Let the number of diploid species be *d* and the number of allotetraploid species be *m*. The network is denoted by *W* and the multi-labeled species tree derived from it is *M*_*W*_. For a given network state, let *h* be the number of hybridizations. Suppose the *i*=1 *i*th allotetraploid subtree has *m*_*i*_ tips (1 ≤ *i* ≤ *h*). Then 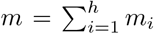. The population size parameters are denoted by the vector *θ*. The parameter *η* is a scaling factor for the population sizes, appearing in a hyperprior for *θ*. The number of gene trees is denoted by *G*. The topology and set of node times for the *i*th gene tree is denoted by *τ*_*i*_ (1 ≤ *i* ≤ *G*). All the other parameters belonging to the *i*th gene tree are denoted by *α*_*i*_; these are parameters for site rate heterogeneity, substitution model, branch rate model, and root model. Thus (*τ*_*i*_, *α*_*i*_) gives all the parameters for the *i*th gene tree. The permutations of sequences within polyploid individuals for the *i*th gene is denoted by *γ*_*i*_. This parameter is the main addition to the usual formula for the multispecies coalescent. We only deal with tetraploids here, so *γ*_*i*_ consists of transpositions (‘flips’) of two sequences. The sequence data for the *i*th gene is denoted by *y*_*i*_. We set *τ* = (*τ*_1_, *…τ*_*G*_), and similarly for *α, γ, y*.

### 2.1 Model

The formula for the posterior density for the AlloppNET model is similar to that used in Jones et al. (2013) and is given by

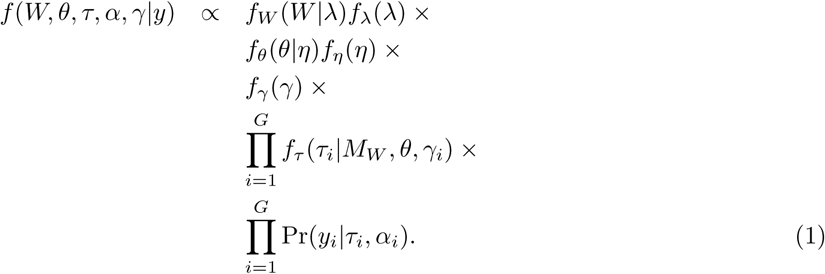

The network prior is *f*_*W*_ (*W |λ*)*f*_*λ*_(*λ*). The choice of this prior poses some problems and is described in a separate section.

The population size prior *f*_*θ*_(*θ| η*)*f*_*η*_(*η*) is for the vector of population size parameters *θ*. There is a mapping from *θ* to values at nodes in the network, and to just after each hybridization event. There are 3*d* + 4*m* + *h* − 2 parameters; note that *h* varies. The population sizes are assumed to vary linearly along edges in the network, except that a instantaneous change is allowed at hybridization events. In the analyses done in this paper, the priors for *θ* used were similar to those typically used by *BEAST. An independent gamma distribution is assumed for each component of *θ*. The shape parameter is 4 for the populations at the tips, 1 for just after hybridizations, and 2 for the rest. The scale parameter for all these gamma distributions is the hyperparameter *η*. The hyperprior *f*_*η*_ for *η* is described later.

The permutation prior *f*_*γ*_ (*γ*) is a discrete distribution on the set of sequence assignments. This is assumed to be uniform here, and thus could be omitted without affecting the inference.

The term *f*_*τ*_ (*τ*_*i*_|*M*_*W*_, *θ*, *γ*_*i*_) provides the probability of *τ*_*i*_, when permuted by *γ*_*i*_, fitting into the multi-labeled species tree *M*_*W*_ with population sizes determined by *θ*. The value of *γ*_*i*_ determines how the sequences for the *i*th gene are assigned to tips in *M*_*W*_. Note that this probability does not depend on *α*_*i*_. Apart from this extra complexity due to the permutations, the value of *f*_*τ*_ (*τ*_*i*_ |*M*_*W*_, *θ*, *γ*_*i*_) is given by the multispecies coalescent, as used in Rannala and Yang (2003), Heled and Drummond (2010) and elsewhere.

The term Pr(*y*_*i*_|*τ*_*i*_, *α*_*i*_) is the probability of the data for the *i*th gene given the *i*th gene tree and other parameters *α*_*i*_. Regarded as a likelihood, it is the usual ‘Felsenstein likelihood’. Here *α*_*i*_ contains the substitution model parameters, branch rate model parameters, and site rate heterogeneity model parameters for the *i*th gene tree. In this paper, we used the HKY substitution model, and assumed strict clock branch rates, and no site rate heterogeneity. The clock rate for one gene was fixed to 1.0, and the others were estimated.

The priors for the population parameter *η* and the parameter *λ* appearing in the network prior, and the priors for relative clock rates were diffuse lognormals.

### Network Prior

There are two difficulties. Firstly, there is very little empirical evidence to guide the choice of prior. Secondly, there is little in the way of theory about probability densities on networks, especially when the number of nodes can vary as it does here. The situation can be contrasted with that of species delimitation (eg Yang and Rannala (2010)) where species trees with different numbers of nodes are considered. In that case, the theory of birth-death processes provides normalized densities for species trees of different sizes which can then be used to determine the Hasting ratios in the MCMC algorithm. But for allopolyploid networks no such densities are known. We therefore resort to writing down a formula for an unnormalized density for each possible size of network, and estimating the properties of the prior by sampling from it.

There are 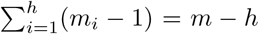 internal nodes in the allotetraploid trees. There are *h* hybridization times, and the diploid history has *d* + 2*h* − 1 internal nodes. The total number of parameters (node times and hybridization times) which are operated on by the reversible jumps is thus *n*:= *d* + *m* + 2*h* − 1. If the ratios between different models (different *h*) is to be the same regardless of *λ*, then the density must reflect this. The formula we use is

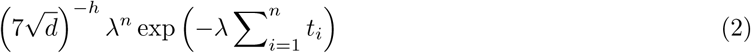

where the *t*_*i*_ are all the node times and hybridization times. The factor 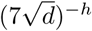 was chosen experimentally so that the marginal distribution over *h ∈*{1*, … …m*} was approximately uniform. Note that *λ* something like a diversification rate.

## 3 MCMC implementation

The network can be represented as a set of allotetraploid subtrees and a ‘diploid history’, as in Figure 2. The diploid history is an ordinary tree with some tips (‘hybridization tips’) having nonzero times. There is one pair of hybridization tips for each hybridization and both tips have the same time, which is the time at which hybridization occurred and hence also the time of origin for an allotetraploid subtree. We will also refer to the external branches in the diploid history which lead to the hybridization tips as the ‘legs’ of the corresponding allotetraploid subtree.

**Figure 2:**
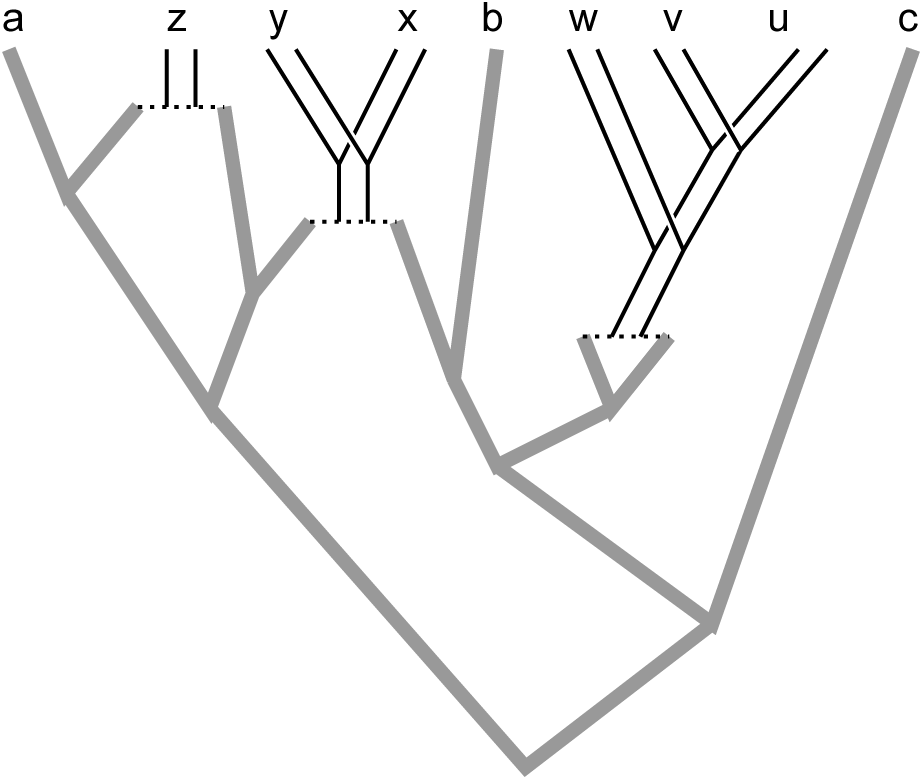
Network with three allotetraploid subtrees (pairs of thin black lines) and the diploid history (thick gray lines). The allotetraploid subtrees have sets {z}, {x,y}, and {w,v,u} of extant species. The diploid history has extant species a,b, and c, plus three pairs of tips which each lead to an allotetraploid subtree.

In this section we describe the MCMC operators (moves) used, and the choice of initial state. There are five novel types of move which are particular to allopolyploid networks:

1. Change a hybridization time.
2. Change an allotetraploid subtree, tipwards of the hybridization
3. Change the diploid history, rootwards of hybridizations.
4. Change the number of allotetraploid subtrees, that is, the number of hybridizations.
5. Change the assignment of sequences within polyploid individuals.

The first is straightforward. The next two have much in common and are described next.

### 3.1 Allotetraploid subtrees and diploid history

Like *BEAST, we use a MCMC move for the species tree based on the ideas of Mau et al. (1999). One reason for using this move is that it can be constrained to keep the species tree compatible with the gene trees. This MCMC move randomly assigns ‘left’ and ‘right’ labels to the immediate descendants of each node, to produce an oriented tree Gernhard (2008) and then alters a node height. In our situation this must take into account the assignment of sequences within allotetraploid individuals. Otherwise, within a single allotetraploid subtree the situation is very similar to that in *BEAST.

The same type of move can be adapted to deal with the diploid history. There are three types of constraint on the new height. Firstly, there are constraints from the gene trees as in the tetraploid subtrees, but the calculation is more complex. In order to calculate the sets of sequences belonging to the left and right subtrees (in the MUL-tree) of a particular node in the diploid history, it is necessary to visit the tetraploid subtrees which are attached to the hybridization tips of the diploid history. Secondly, there are lower bounds on the new height due to the fact that the hybridization tips have nonzero height. In the oriented tree, this amounts to ensuring that the new height does not become smaller than either of the heights of adjacent nodes. (Nodes adjacent to internal nodes are always tips in the left-right ordering.) Thirdly there are constraints to keep the root as a diploid. If node to change height is the root, and the second highest node is to left or right of all diploids, then the root must stay the root: there is a lower limit which is the height of second highest node. If the node to slide is not the root, and is to left or right of all diploids, then it must not become the root: there is an upper limit which is the root height.

### 3.2 The number of hybridizations

This is the most complex move. Sampling all values of *h* can be done by repeatedly changing *h* to *h* − 1 or *h* + 1, and that can be done by splitting one tetraploid subtree into two and merging two into one. The difficult part is making the moves reversible, so that the probability of a move going from one network state A to another B is balanced by a reverse move. When the MCMC move for changing *h* is chosen, a split or a merge is chosen with equal probability. If a split is chosen, but no splits are possible, no move is made; the same network state is sampled again. Likewise, if a merge is chosen, but no merges are possible, the same network state is sampled again.

Splitting (going left to right in the top section of Fig 3). Any tetraploid subtree with more than one tip can be split. One, T, is chosen at random. The two child nodes of the root of T become the roots of the two new tetraploid subtrees. The child nodes are not treated symmetrically in the move, so both orderings of the child nodes is treated as candidates. There are thus twice as many candidate splits as tetraploid subtrees. The steps are:

1. Split T into T1 and T2 and create a new hybridization height for T1 between the root height of T1 and the root height of T.
2. Create a new hybridization height for T2 between the root height of T2 and the root height of T. Create two ancestor nodes for the hybridization tips, one re-using the root height of T and the other between this time and the minimum of the limits imposed by the gene trees and the height of the node that will become its ancestor node.
3. Join up the topology in the diploid history. Note that they could join the diploid history in many ways but a particular way is always chosen, so that the two new subtrees ‘share legs’ as shown. Note also that it is necessary to keep track of which of the two ‘copies’ of a tetraploid subtree is which (ie, the left leg of one must correspond to the left leg of the other).

The most difficult parameter is the new node height when splitting, which is below the root height of the subtree. This can conflict with gene trees, so the sets of (species,sequence) pairs have to be found for the two child nodes of the new node, and the gene trees examined to find the most recent coalescence that conflicts with the new node.

**Figure 3:**
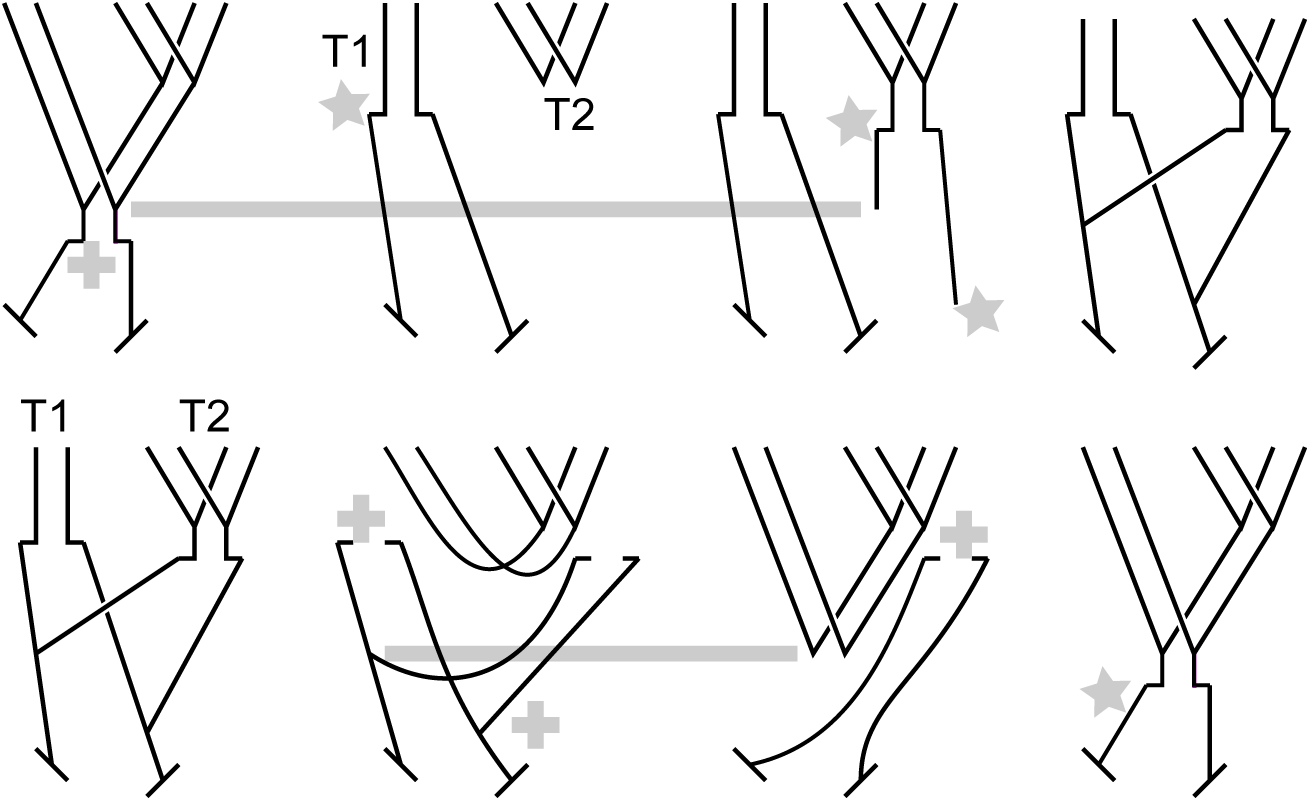
MCMC moves to change the number of tetraploid subtrees. Top: splitting. Bottom: merging. The two gray horizontal lines show times that are re-used. Gray crosses indicate times that are about to disappear. Gray stars show times which have just been created.

Merging (going right to left in the bottom section of Fig 3) must be the reverse of splitting. So two tetraploid subtrees can only merge if they have a configuration like that in the figure, ie ‘sharing legs’. It is not necessary that the two nodes at the bottom of the figure be different. A list of possible pairs of tetraploid subtrees is made, and if there any suitable pairs, one pair T1,T2, is chosen at random, and the merge is carried out. The steps are:

1. Merge T1 and T2 into T. The root height of T becomes the height of the most recent ancestor to a hybridization tip.
2. Remove the hybridization tips for T1 from the diploid history. This loses a hybridization height and a node height. The latter requires finding the limit from gene trees for Hastings ratio.
3. Give the hybridization tips for T1 new heights below the root height of T, and join up.

Note that the limit from gene trees must be calculated for the lost node height (as when splitting), in order to calculate the Hastings ratio.

We use the theory developed in Green (1995) to calculate the Hastings ratios for splitting and merging moves. Consider a splitting move; the merging case is similar. Then equation (7) of Green (1995) provides the acceptance ratio

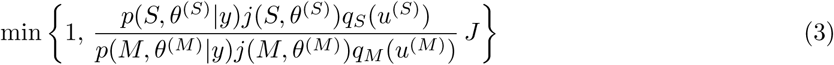

where

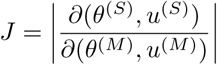

is the Jacobian and *S* (for ‘split’) and *M* (for ‘merged’) replace Green’s 1 and 2. Here *y* is the data, and *p*(*S, θ*^(*S*)^*|y*) and *p*(*M, θ*^(*M*)^*|y*) the usual Bayesian posteriors. The term *j*(*S, θ*^(*S*)^) gives the probability of choosing the splitting move and *j*(*M, θ*^(*M*)^) of choosing the reverse merging move. The term *θ*^(*S*)^ is a vector of node and hybridization times for the network with the split case, and *θ*^(*M*)^ is a vector of node and hybridization times for the network with the merged case. The vectors *u*^(*S*)^ of length 3 and *u*^(*M*)^ of length 1 provide the extra parameters created when doing the jump. The function *q*_*S*_ is the density of the distribution from which *u*^(*S*)^ is sampled; likewise *q*_*M*_.

In our case *u*^(*S*)^ and *u*^(*M*)^ can be generated by independent sampling from the uniform distribution on [0, 1] for each dimension. In this case *q*_*S*_ (*u*^(*S*)^) and *q*_*M*_ (*u*^(*M*)^) are both 1 and can be omitted from the formula. The new parameters are then derived from these values, as functions of the other old parameters. In the present case, all these function are linear functions mapping [0, 1] to a suitable range, the range being some function of the other parameters *θ*^(*S*)^ or *θ*^(*M*)^. The probabilities of the moves being chosen, namely *j*(M*, θ*^(*M*)^) and *j*(S*, θ*^(*S*)^) must also be taken into account.

When the number of hybridizations changes, the number of population parameters also changes. It increases by one for each hybridization. So in a split a new population parameter is added, and in a merge one is removed. When one is added, it is sampled from the prior for the population. The contribution to the Hastings ratio is calculated from the value of the density of the population prior at the new parameter value when splitting, or at the lost value when merging.

### 3.3 Assignment of sequences within allotetraploid individuals

A uniform prior on the possible assignments is used, so the Hastings ratios are all 1. The reassignment moves need to visit all possible ‘flips’ for each gene in each tetraploid individual. This is easy to arrange in a mathematical sense. The only difficulty is choosing combinations of flips which have good mixing properties. As usual with MCMC algorithms, this requires experimentation.

Three types of move have been implemented. The first flips the assignment of a single gene in a single individual. The second works within a single gene tree, and chooses a random node within it, and roughly speaking, flips a clade of individuals. The details are in the code.

The third type of move was introduced to avoid the MCMC chain getting stuck in a particular situation. The merging move which reduces the number of hybridizations by one, requires the left legs to be ‘shared’, and the right legs to be shared; left leg shared with right leg will not do. However the network can get into a situation where the legs are ‘reversed’ *and* the sequence assignments of the genes at the tips of the two candidate are opposite to one another. This makes a state which may be very difficult to leave. The third move flips sequence assignments of all genes of all individuals of all species in a tetraploid subtree, *and* switches the legs around, swapping left and right. This takes the network to an equivalent state with the same likelihood, but one from which the merging move can operate.

### 3.4 Initial state

A random initial state for the species network is chosen as follows.

1. The tetraploid species are partitioned into one or more groups using the Chinese restaurant process (Kingman, 1993; Pitman, 2006).
2. Trees from a Yule process are generated for each of the groups of tetraploid species.
3. The diploid history is constructed in a manner similar to the Yule process working backwards in time, with some modifications. Each diploid species is a tip and there are two hybridization tips for each tetraploid subtree. Two subtrees are repeatedly chosen from those available and joined into a subtree. When two nodes are selected for joining, the choice is constrained so that the diploids do not all merge while there are still tetraploids left to merge. Also, the height of the root of the new subtree has to be made earlier than either of the nodes chosen for joining (which would not happen automatically, since the hybridization tips have nonzero height).

## 4 Simulations

Three sets of simulations were done.

### 4.1 Sampling from prior

Firstly, the program was run with no data in order to assess the prior on the species network. Various numbers of diploid species *d* and tetraploid species *m* were tested. Four cases with of *d* = 2, 8 and *m* = 2, 8 are reported here.

### 4.2 Scenarios D,E,F

Secondly, a large number of simulations were run for three scenarios labeled D,E, and F. They are shown in Figure 4. The three topologies were chosen to be similar but have different numbers of hybridizations. Different numbers of genes (*G* = 1, 3, 9), individuals per species (*N* = 1, 3, 9), and mutation rates (*T* = 4e-9, 2e-8, 1e-7 mutations per site per generation) were tested. For each of the 81 possible values, ten replicates were generated and analyzed.

All genes were 500bp long. Population sizes were 100,000 individuals (hence 200,000 gene copies per diploid genome) at the tips, and at rootward ends of branches, and 200,000 individuals at tipward ends of internal branches and at the root. Strict clock branch rates, no site rate heterogeneity, and equal clock rates for all genes were assumed when simulating the data. The HKY substitution model was assumed and parameter kappa was set to 3, and the frequencies set to .3 for A and T, and .2 for C and G (Seq-Gen was called with parameters -t3.0 -f0.3,0.2,0.2,0.3). These substitution parameters were estimated in the analysis. Priors on population size scaling factor *η*, the relative mutation rates of genes, and *λ* were all diffuse log-normals.

Note that since the root height is kept the same in terms of substitutions when the mutation rate is varied, different values of the mutation rate mean different numbers of generations from root to tip. Large values of *T* result in greater amounts of incomplete lineage sorting. When *T* = 4e-9, the root height measured in number of generations is 3,000,000. For *T* = 2e-8 it is 600,000 and for *T* = 1e-7 it is 120,000.

### 4.3 Scenario J

The third set of simulations used a single, more complex scenario with 6 diploid species and 7 allotetraploid species, as shown in Figures 5 and 6. The settings for gene lengths, population sizes, site model parameters, substitution model parameters, and priors were the sames as for scenarios D, E, and F. Only one value of *T*, namely 2e-8, was used. Values of *G*=1,3,9 and *N* =1,3 were used. Ten replicates were run for each case.

**Figure 4:**
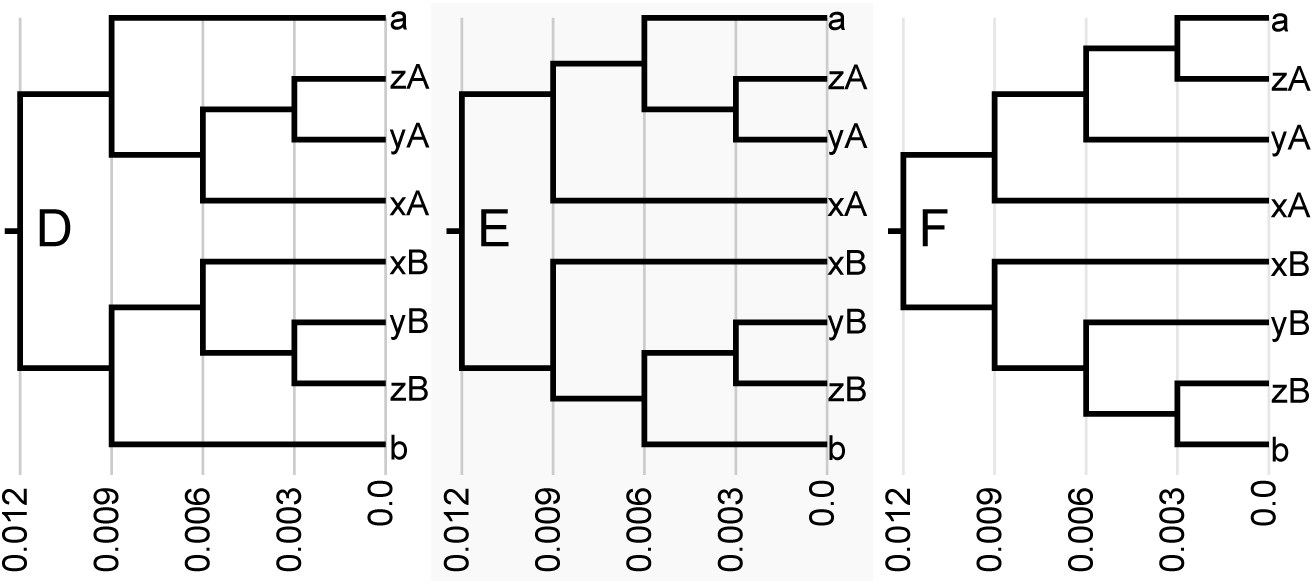
Scenarios D,E,F: the true MUL-trees. D has one hybridization; E has two; and F has three. Heights are in expected numbers of substitutions.

**Figure 5:**
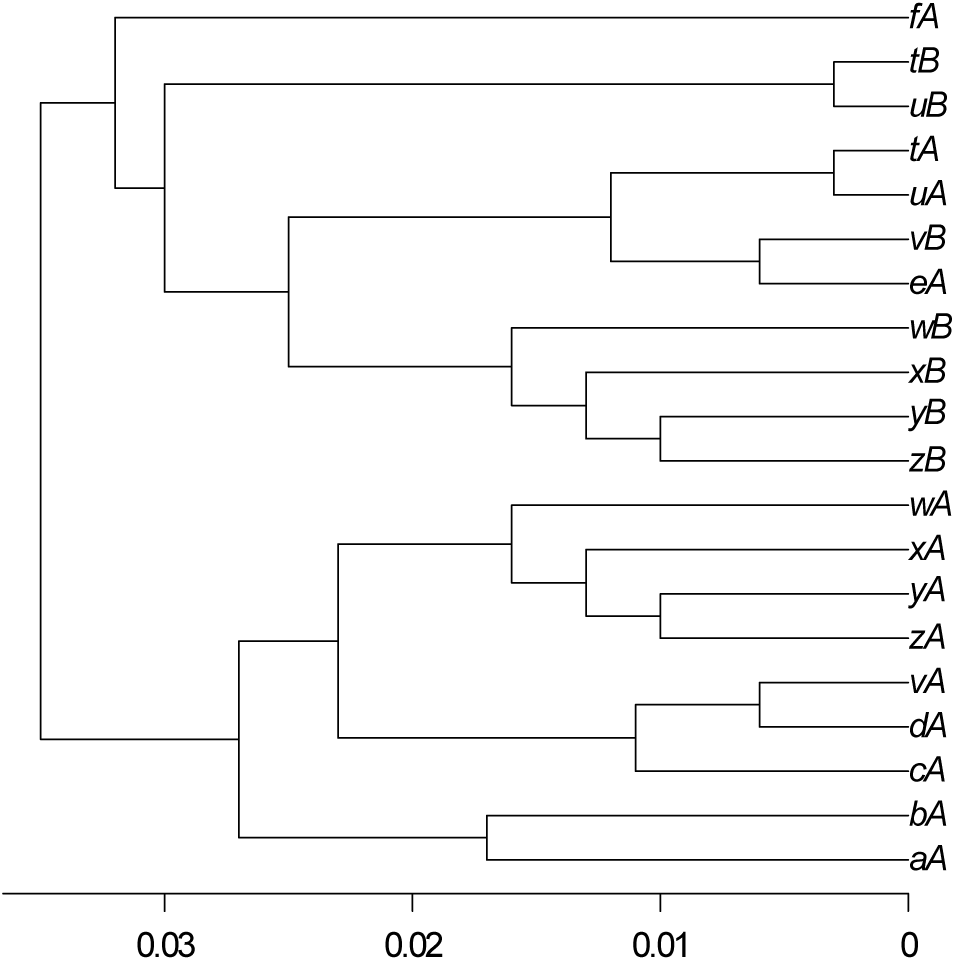
The true MUL-tree for scenario J with 6 diploid species labelled a,b,c,d,e,f and 7 tetraploid species comprising clades {t,u}, {v}, and {w,x,y,z}, arising from 3 hybridizations. Heights are in expected numbers of substitutions.

**Figure 6:**
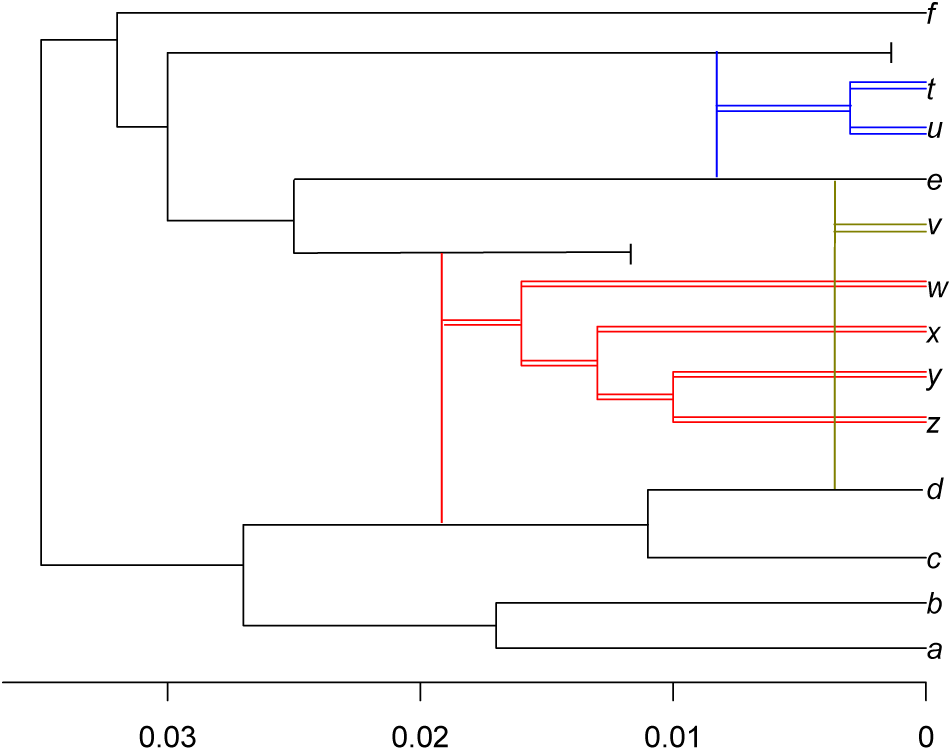
The network for scenario J, corresponding to Figure 5. The colours red, green, and blue indicate the three allotetraploid subtrees. Two diploid species have become extinct.

## 5 Results

### 5.1 Distances for multi-labeled trees

In order to summarize the accuracy of the results it useful to have some definitions of distances for multi-labeled trees. We start with a version of the Robinson-Foulds distance (Robinson and Foulds, 1981) which includes branch lengths and is adapted for binary rooted trees. We denote it by *D*_*RF*_. It is defined by the following algorithm. Given two binary rooted trees *T*_1_ and *T*_2_ with the same tip labels:

1. For each node *i* in *T*_*j*_ (*j* ∈ {1, 2}), find the clade *C*_*ji*_ and the length of the branch *B*_*ji*_ leading to *C*_*ji*_.
2. *D*_*RF*_ = 0.
3. For each clade *C*_*ji*_ which does not have a match in the other tree, add *B*_*ji*_ to *D*_*RF*_.
4. For each clade *C*_*ji*_ which does have a match *C*_*kl*_ (for some *l* and where *k* = 2 if *j* = 1 and *k* = 1 if *j* = 2) in the other tree, add |*B*_*ji*_ - *C*_*kl*_| to *D*_*RF*_.

In order to extend this to multi-labeled trees *M*_1_ and *M*_2_, we follow Huber et al. (2011) and define a distance *D*_*RF* *M*_ by considering all possible consistent relabellings of *M*_1_ and *M*_2_ and finding the minimum distance over all such relabellings. For the results here, this amounts to giving each pair of tips in *M*_1_ an arbitrary labeling to distinguish them (say ‘A’ and ‘B’), and then labeling each pair of tips in *M*_2_ with either (‘A’,‘B’) or (‘B’,‘A’). Thus, in order to evaluate a distance for a pair multi-labeled trees with *m* allotetraploids, 2^*m*^ ordinary Robinson-Foulds-type distances must be evaluated.

### 5.2 Sampling from prior

With no data, the prior given by equation 2 is sampled. Some examples of the marginal distribution for *m* are shown in Figure 7. It is fairly uniform over *m*. This is the case for the range of numbers of diploid and tetraploid species considered here, but not for much larger numbers of species.

**Figure 7:**
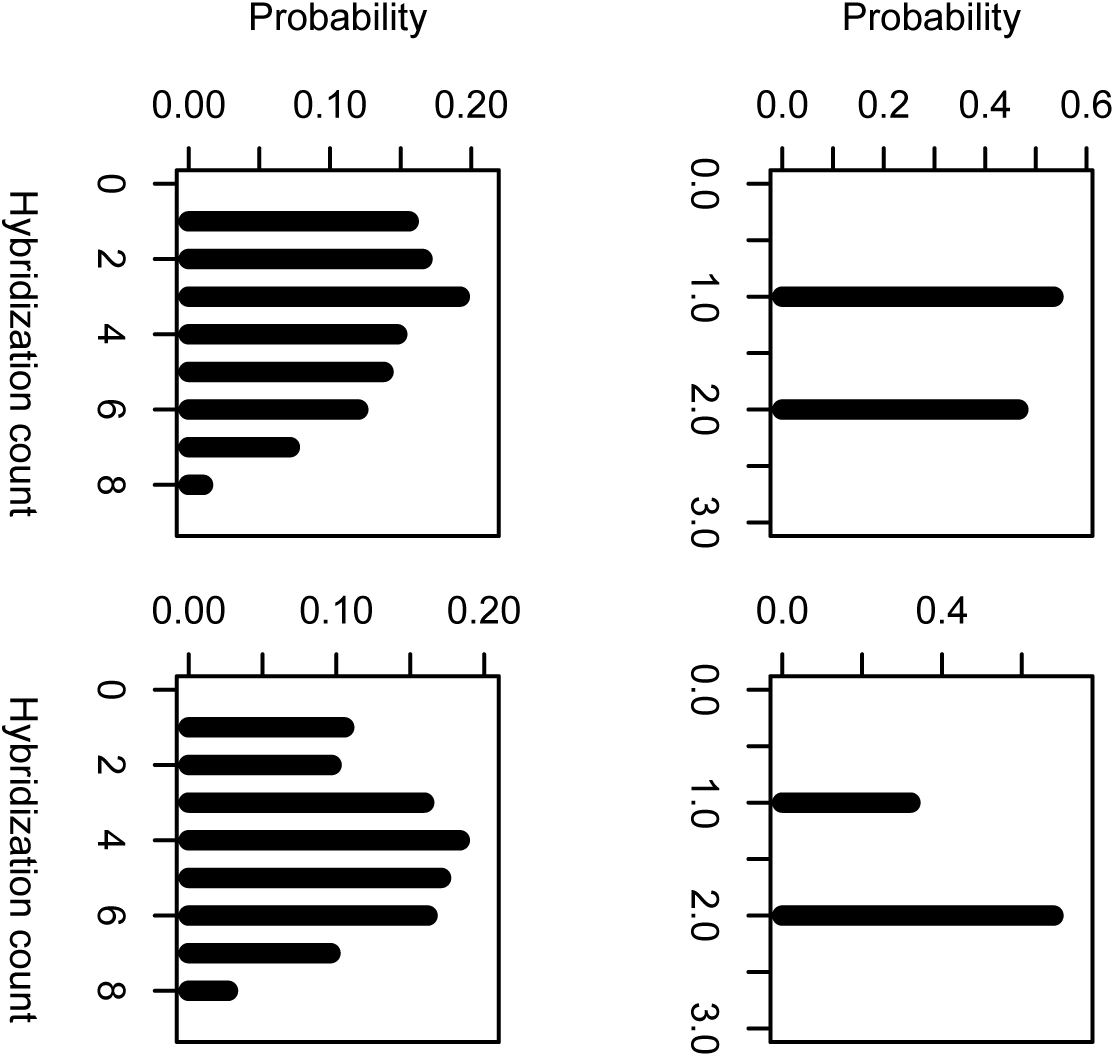
Estimates of the marginal distribution for the number of hybridizations *m* in the prior. There are 2 tetraploid species in the top row, and 8 in the bottom two. There are 2 diploid species in the left column and 8 in the right column.

## 5.3 Scenarios D, E, and F

MCMC chains of 30M generations were used. Burnin was set to remove the first 5M. Note that although there are only 5 species, there are 8 diploid genomes and therefore 8 *×* 9 *×* 9 = 648 sequences for the *N* = 9*, G* = 9 case. The BEAST runs for this case took around 4 hours each using one core on a desktop computer.

For each of the 27 sets of *G, N, T* values, the *D*_*RF* *M*_ values were similar for the three scenarios and were combined to produce the results shown in Figure 8. As expected, accuracy increases with *G*, *N*, and decreases with *T*. In general, increasing *G* is more useful than increasing *N*. It is worth noting that estimates of *m* are often correct even when the topology is wrong (results not shown).

## 5.4 Scenario J

MCMC chains of 30M generations with burnin set to remove the first 5M were used for most *G* and *N* values, but 100M generations with burnin set to remove the first 10M was used for *G* = 9*, N* = 3. The results are shown in Figure 9.

**Figure 8:**
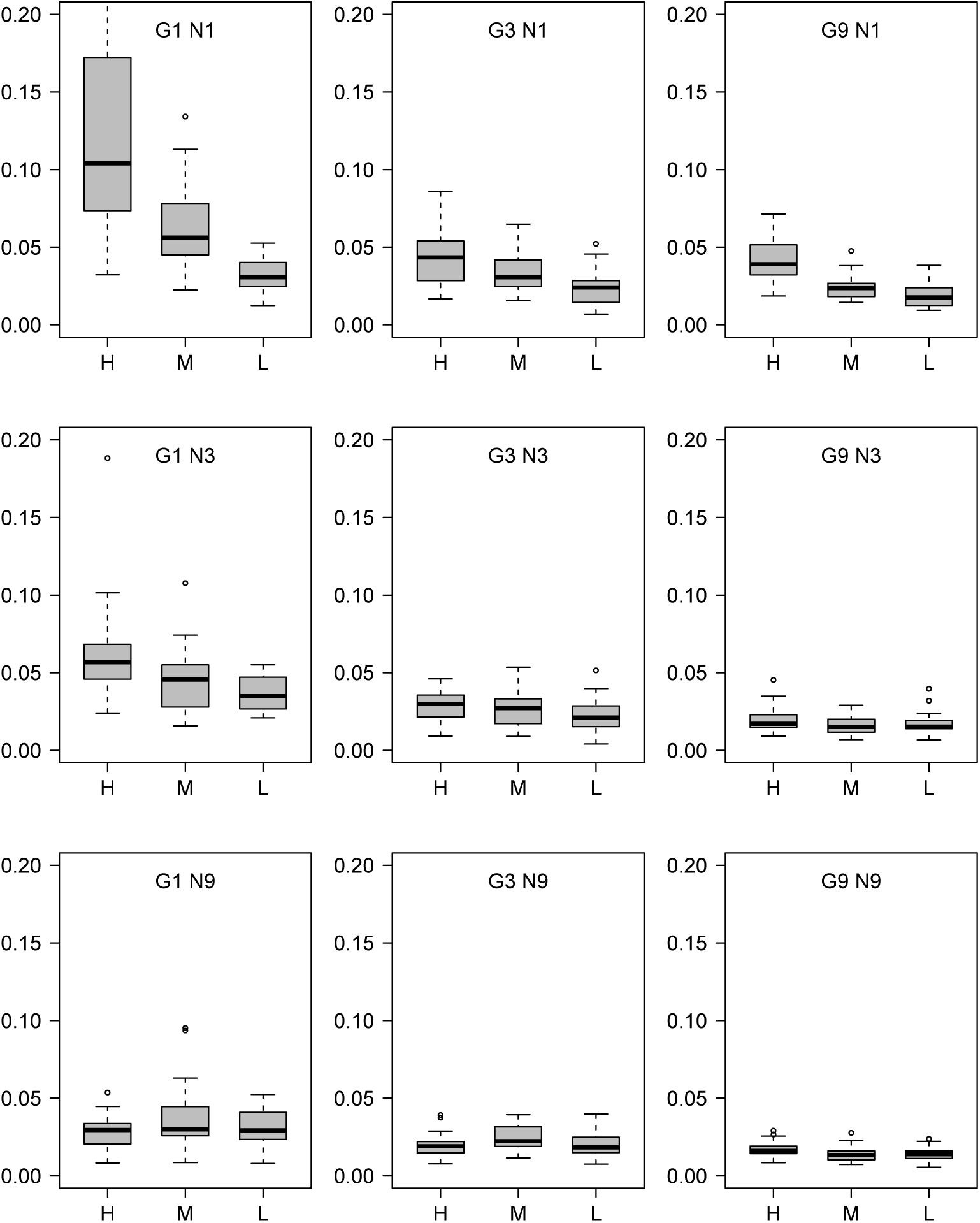
Results for scenarios D,E,F combined. The boxplots show values over 10 replicates per scenario for *D*_*RF* *M*_ values for different values of the number of loci *G* and individuals per species *N*. Each graph shows results for different amounts of incomplete lineage sorting (H = high, M = medium, L = low).

**Figure 9:**
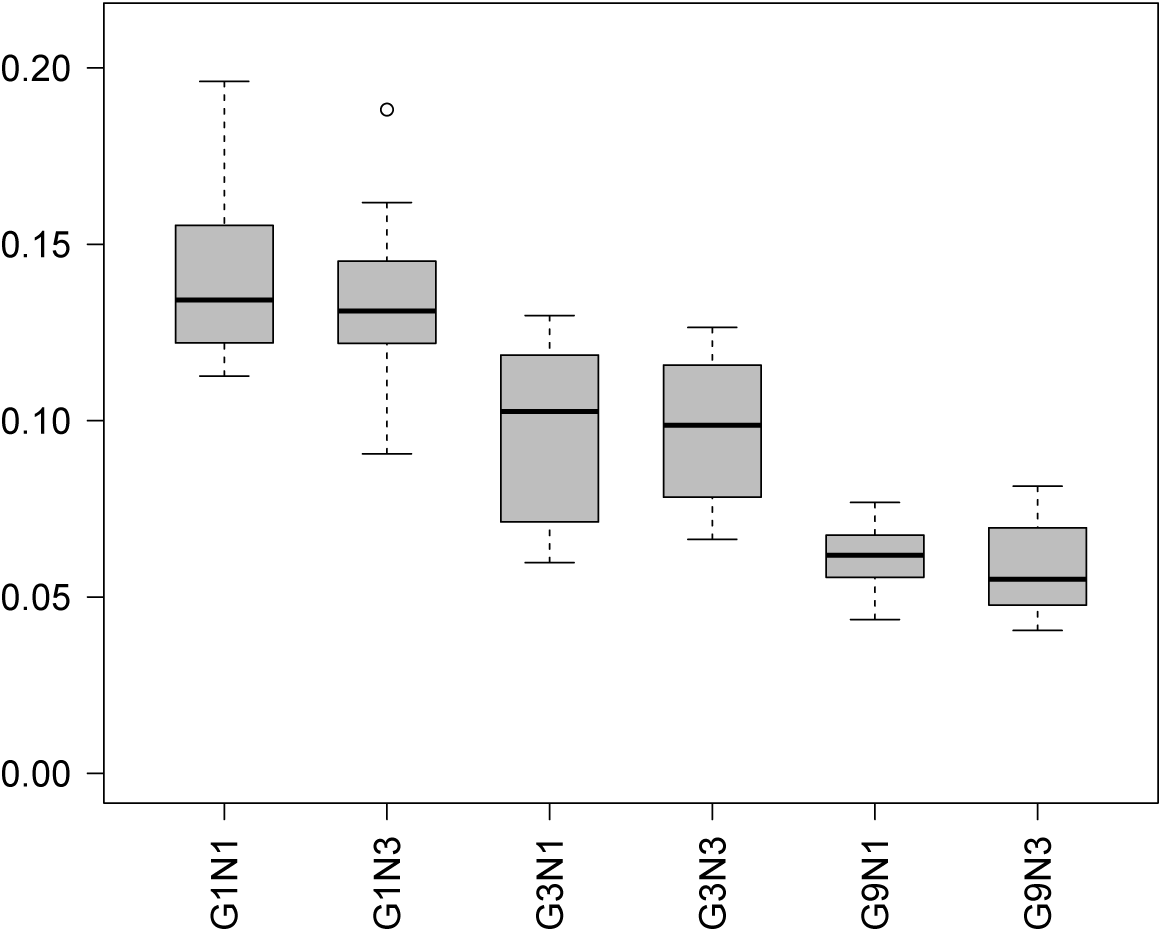
Results for scenario J. The boxplot shows mean values over 10 replicates for *D*_*RF* *M*_ values.

## References

T Gernhard. The conditioned reconstructed process. J. Theo. Biol., 253:769–778, 2008.

Peter J Green. Reversible jump Markov chain Monte Carlo computation and Bayesian model determination. Biometrika, 82(1):711–732, 1995.

J Heled and A Drummond. Bayesian inference of species trees from multilocus data. Mol. Biol. Evol., 27: 570–580, 2010.

K T Huber and V Moulton. Phylogenetic networks from multilabelled trees. J Math Biol, 52:613–632, 2006.

K T Huber, A Spillner, R Suchecki, and V Moulton. Metrics on multi-labelled trees: interrelationships and diameter bounds. IEEE/ACM Transactions on Computational Biology and Bioinformatics, 8(4):1029–1040, 2011.

G Jones, S Sagitov, and B Oxelman. Statistical inference of allopolyploid species networks in the presence of incomplete lineage sorting. Syst. Biol., 2013. doi: 10.1093/sysbio/syt012.

J.F.C. Kingman. Poisson Processes. Oxford University Press, 1993.

Bob Mau, Michael A Newton, and Bret Larget. Bayesian phylogenetic inference via Markov chain Monte Carlo methods. Biometrics, 55:1–12, 1999.

J. Pitman. Combinatorial Stochastic Processes. Ecole d’été de Probabilités de St-Flour XXXII, Lecture Notes in Mathematics 1875. Springer, 2006.

B Rannala and Z Yang. Bayes estimation of species divergence times and ancestral population sizes using DNA sequences from multiple loci. Genetics, 164:1645–1656, 2003.

D. Robinson and L. Foulds. Comparison of phylogenetic trees. Math. Biosciences, 53:131–147, 1981.

C J Rothfels, K M Pryer, and F W Li. Next generation polyploid phylogenetics: rapid resolution of hybrid polyploid complexes using pacbio single-molecule sequencing. New Phytol., 213:413–429, 2017.

Z Yang and B Rannala. Bayesian species delimitation using multilocus sequence data. Proceedings of the National Academy of Sciences of U.S.A, 107:9264–9269, 2010.

